# Hydroxyl carlactone derivatives are predominant strigolactones in *Arabidopsis*

**DOI:** 10.1101/2020.01.17.910877

**Authors:** Kaori Yoneyama, Kohki Akiyama, Philip B. Brewer, Narumi Mori, Miyuki Kawada, Shinsuke Haruta, Hisashi Nishiwaki, Satoshi Yamauchi, Xiaonan Xie, Mikihisa Umehara, Christine A. Beveridge, Koichi Yoneyama, Takahito Nomura

**Affiliations:** Graduate School of Agriculture, Ehime University, Matsuyama 790-8566, Japan; PRESTO, Japan Science and Technology, Kawaguchi, Saitama 332-0012, Japan; Department of Applied Life Sciences, Graduate School of Life and Environmental Sciences, Osaka Prefecture University, Sakai, Osaka 599-8531, Japan; ARC Centre of Excellence in Plant Energy Biology, School of Agriculture, Food and Wine, The University of Adelaide, Glen Osmond, SA 5064, Australia; Center for Bioscience Research and Education, Utsunomiya University, Utsunomiya 321-8505, Japan; Department of Applied Biosciences, Faculty of Life Sciences, Toyo University, Gunma 374-0193, Japan; ARC Centre of Excellence for Plant Success in Nature and Agriculture, School of Biological Sciences, The University of Queensland, St. Lucia, QLD 4072, Australia; Women’s Future Development Center, Ehime University, Matsuyama 790-8577 Japan

**Keywords:** *Arabidopsis thaliana*, hydroxyl carlactone derivative, lateral branching oxidoreductase

## Abstract

Strigolactones (SLs) regulate important aspects of plant growth and stress responses. Many diverse types of SL occur in plants, but a complete picture of biosynthesis remains unclear. In *Arabidopsis thaliana*, we have demonstrated that MAX1, a cytochrome P450 monooxygenase, converts carlactone (CL) into carlactonoic acid (CLA), and that LBO, a 2-oxoglutarate-dependent dioxygenase, converts methyl carlactonoate (MeCLA) into a metabolite called [MeCLA+16] Da. In the present study, feeding experiments with deuterated MeCLAs revealed that [MeCLA+16] Da is hydroxymethyl carlactonoate (1’-HO-MeCLA). Importantly, this LBO metabolite was detected in plants. Interestingly, other related compounds, methyl 4-hydroxycarlactonoate (4-HO-MeCLA) and methyl 16-hydroxycarlactonoate (16-HO-MeCLA) were also found to accumulate in *lbo* mutants. 3-HO-, 4-HO- and 16-HO-CL were detected in plants, but their expected corresponding metabolites, HO-CLAs, were absent in *max1* mutants. These results suggest that HO-CL derivatives are predominant SLs in *Arabidopsis*, produced through MAX1 and LBO.

## INTRODUCTION

Strigolactones (SLs) were originally identified as germination stimulants for root parasitic plants (Cook et al., 1966) and then as hyphal branching factors for symbiotic arbuscular mycorrhizal (AM) fungi (Akiyama et al., 2005). SLs were thought to function only as rhizosphere signals until the discovery of their role as a plant hormonal signal that inhibits lateral shoot branching (Gomez-Roldan et al., 2008; Umehara et al., 2008).

Shoot branching involves the formation of axillary buds in the axil of leaves. The level of dormancy in buds is an essential determinant of plant architecture. Defects in the SL pathway correspond with loss of bud dormancy and excessive shoot branching as displayed by SL mutants that include *ramosus* (*rms*) of pea (*Pisum sativum*), *decreased apical dominance* (*dad*) of petunia (*Petunia hybrida*), *dwarf* (*d*) of rice (*Oryza sativa*) and *more axillary growth* (*max*) of *Arabidopsis* (*Arabidopsis thaliana*).

Natural SLs are carotenoid-derived compounds consisting of a butenolide D ring linked by an enol ether bridge to a less conserved moiety. These SLs can be classified into two structurally distinct groups: canonical and non-canonical SLs. Canonical SLs contain the ABCD ring formation, and non-canonical SLs lack the A, B, or C ring but have the enol ether-D ring moiety (Al-Babili and Bouwmeester, 2015). During biosynthesis, the initial compound that contains the D ring is carlactone (CL), an endogenous precursor for SLs, which is produced by the sequential reactions of 9-*cis*/all-*trans*-β-carotene isomerase and two carotenoid cleavage dioxygenases (CCD7, CCD8) (Alder et al., 2012). In *Arabidopsis*, the isomerase is encoded by *DWARF27* (*D27*), and CCD7 and CCD8 by *MAX3* and *MAX4*, respectively (Fig. 1). We have demonstrated that recombinant MAX1, (a cytochrome P450 monooxygenase) expressed in yeast, converts CL to carlactonoic acid (CLA) by oxidations at C-19 (Abe et al. 2015). This function was also observed in MAX1 homologs of other plant species including rice, maize, tomato, a model tree poplar, and a lycophyte spike moss, suggesting this conversion of CL to CLA is highly conserved in the plant kingdom (Yoneyama et al., 2018). It was also shown that CL, CLA, and methyl carlactonoate (MeCLA) are present in *Arabidopsis* root tissues (Seto at al., 2014; Abe et al., 2014). Furthermore, differential scanning fluorimetry and hydrolysis activity tests showed that, among CL, CLA, and MeCLA, only MeCLA could interact with the SL receptor, AtD14, suggesting MeCLA may be biologically active in the inhibition of shoot branching in *Arabidopsis* (Abe et al., 2015). *Arabidopsis max1* mutants display a highly increased lateral shoot branching phenotype, and yet accumulate CL (Seto et al., 2014), indicating that CL is not active in repressing shoot branching.

**Figure 1.**
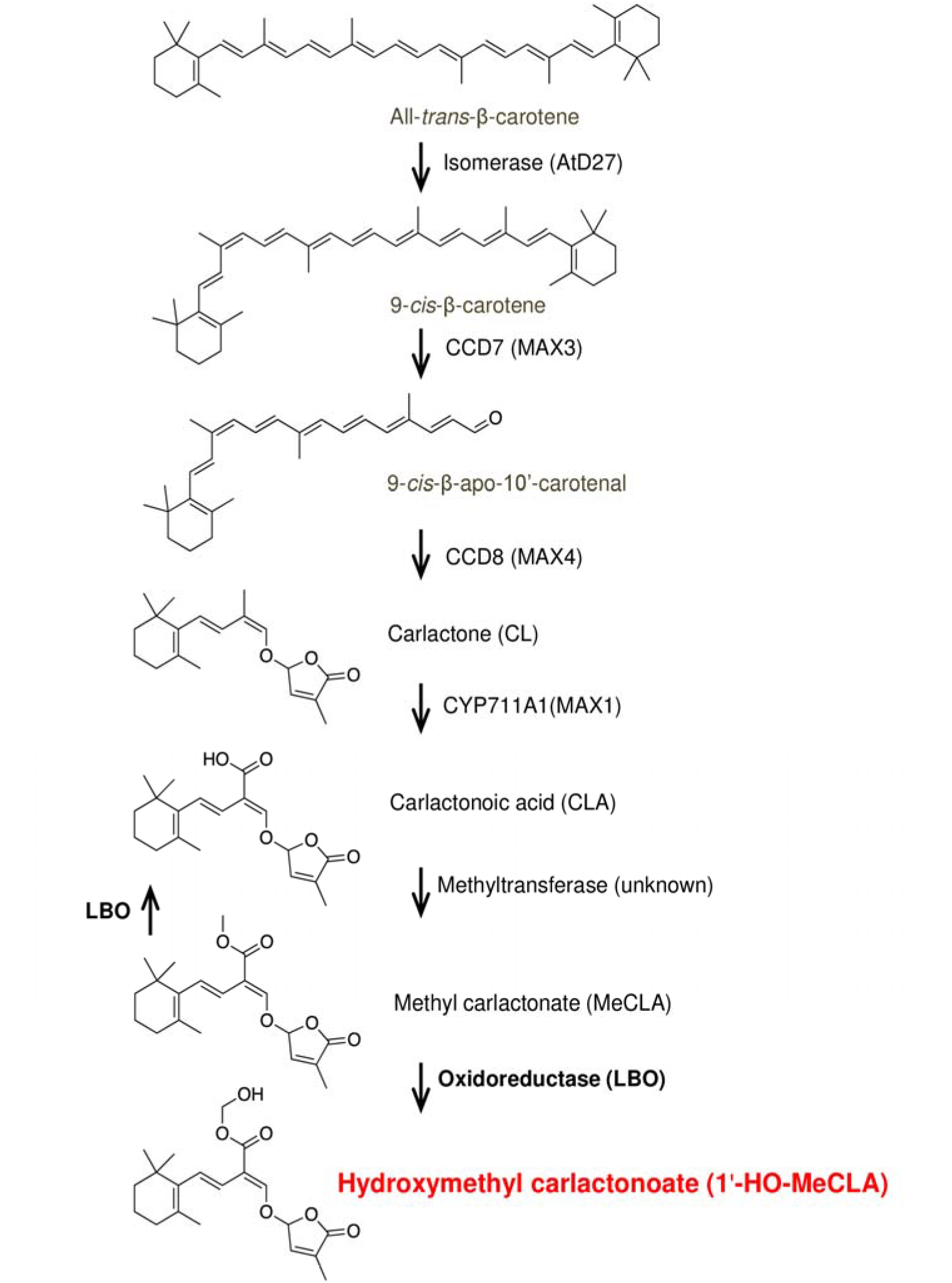
Proposed strigolactone (SL) biosynthesis pathway in *Arabidopsis thaliana.* An isomerase (AtD27) and two CCD enzymes (MAX3 and MAX4) convert β-carotene into carlactone (CL), an endogenous common precursor for diverse SLs. CL is then oxidized by cytochrome P450 (MAX1) to carlactonoic acid (CLA), which is converted into MeCLA by unknown methyltransferase. The present study showed that 2-oxoglutarate-dependent dioxygenase LBO converts MeCLA into 1’-HO-MeCLA, which is essential for regulating shoot branching.

As a novel SL biosynthetic gene, *LATERAL BRANCHING OXIDOREDUCTASE* (*LBO*), encoding a 2-oxoglutarate and Fe (II)-dependent dioxygenase was identified by using a transcriptomic approach, and was shown to function downstream of MAX1 (Brewer et al., 2016). *Arabidopsis lbo* mutant shoot branching is increased compared to WT (Ws-4), but its phenotype is intermediate between WT and *max4* mutants. LC-MS/MS analysis of SLs revealed that CL and MeCLA accumulate in root tissues of *lbo* mutants (Brewer et al., 2016). Because the active shoot branching inhibitor MeCLA accumulates in *lbo* mutants, the intermediate branching phenotype of *lbo* mutants may be explained by the presence of MeCLA. Thus, it was suggested that LBO is necessary for complete suppression of shoot branching in plants by converting the partly bioactive MeCLA to a compound with greater bioactivity for branching. We then showed that the LBO enzyme expressed in *E. coli* only consumed MeCLA when fed with CL, CLA, or MeCLA, and converted MeCLA into a product of [MeCLA+16] Da. However, complete characterization of this LBO metabolite had not yet been conducted.

In the present study, we have determined the structure of the [MeCLA+16] Da compound produced by LBO from MeCLA by feeding experiments using deuterated MeCLAs. In addition, we could identify this LBO metabolite as an endogenous compound from not only roots, but also basal parts of *Arabidopsis* shoot tissues. Since two additional *lbo* mutant alleles, *lbo-2* and *lbo-3,* exist, and homozygous mutant plants exhibited increased shoot branching (Brewer et al., 2016), recombinant proteins of LBO-2 and LBO-3 were produced and the correlation between their enzymatic activities in the conversion of MeCLA to [MeCLA+16] Da and their shoot branching phenotypes was investigated to further examine the importance of the LBO metabolite for shoot branching. Then, biochemical functions of LBO homologs in other plant species including tomato, maize, and sorghum were examined to clarify if the conversion of MeCLA to [MeCLA+16] Da is conserved among these plant species. Furthermore, endogenous SLs in *Arabidopsis max1* and *lbo* mutants were carefully analyzed in search of other potential substrates for MAX1 and LBO to better understand the SL biosynthetic pathway in *Arabidopsis*.

## RESULTS

### LBO catalyzes the conversion of methyl carlactonoate (MeCLA) into hydroxymethyl carlactonoate (1’-HO-MeCLA)

To characterize the structure of [MeCLA+16] Da, LBO enzyme reactions were performed repeatedly. Both the substrate MeCLA and the metabolite [MeCLA+16] Da were highly unstable and the yield of the metabolite was extremely low. We tried to optimize enzyme assay conditions but the maximum yield of the LBO metabolite did not exceed 0.1%. Although more than 500 μg of synthetic MeCLA has been used for LBO enzyme assay, the amount of the metabolite after purification by DEA, silica, and HPLC was not enough for NMR spectroscopy measurement.

The observed mass of [MeCLA+16] Da (Brewer et al. 2016) suggests that LBO has simply added an oxygen to MeCLA. Therefore, feeding experiments with using deuterated MeCLAs were conducted to identify the site of oxidation of MeCLA (Nomura et al., 2013). When MeCLA was fed to LBO, the metabolite was detected by the transition of *m/z* 363 to 97 (Fig. 2). When 18-*d_3_*-MeCLA was fed, the metabolite was detected by the transition of *m/z* 366 to 97 (Fig. 2), clearly indicating that 18-*d_3_* remained unaffected and thus oxidation did not occur at C-18. By contrast, when 1’-*d_3_*-MeCLA, in which ester methyl group had been labeled with deuterium was fed, major metabolite was detected by the transition of *m/z* 365 to 97 (Fig. 2), apparently showing that ester methyl group was oxidized. Consequently, it was demonstrated that LBO converts MeCLA into hydroxymethyl carlactonoate (1’-HO-MeCLA) (Fig. 1).

**Figure 2.**
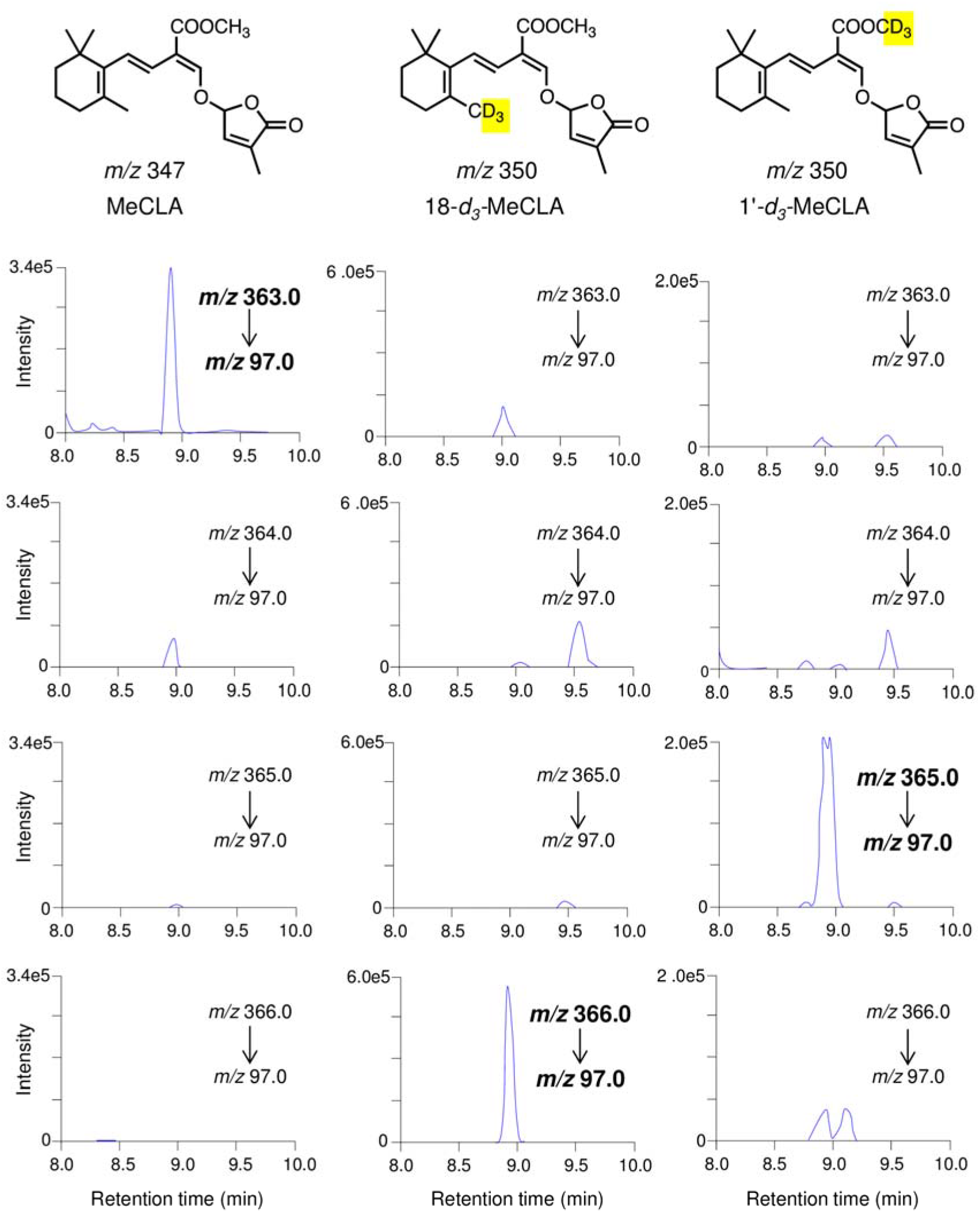
LBO converted [18-*d*_3_]-MeCLA to [MeCLA+16+3] and [1’-*d*_3_]-MeCLA to [MeCLA+16+2]. To characterize the structure of [MeCLA+16], [18-*d*_3_]-MeCLA (*Middle*) and [1’-*d*_3_] MeCLA (*Light*) were fed as substrates to recombinant LBO proteins and incubated for 15 min. Products were identified by LC-MS/MS (MRM).

On the other hand, when MeCLA was incubated with LBO, most MeCLA was converted to CLA; the ratio of CLA to 1’-HO-MeCLA was 100: 1 based on the peak areas in the LC-MS/MS chromatograms of LBO reaction products (Fig. 3), indicating that the LBO protein assay mainly produces CLA.

**Figure 3.**
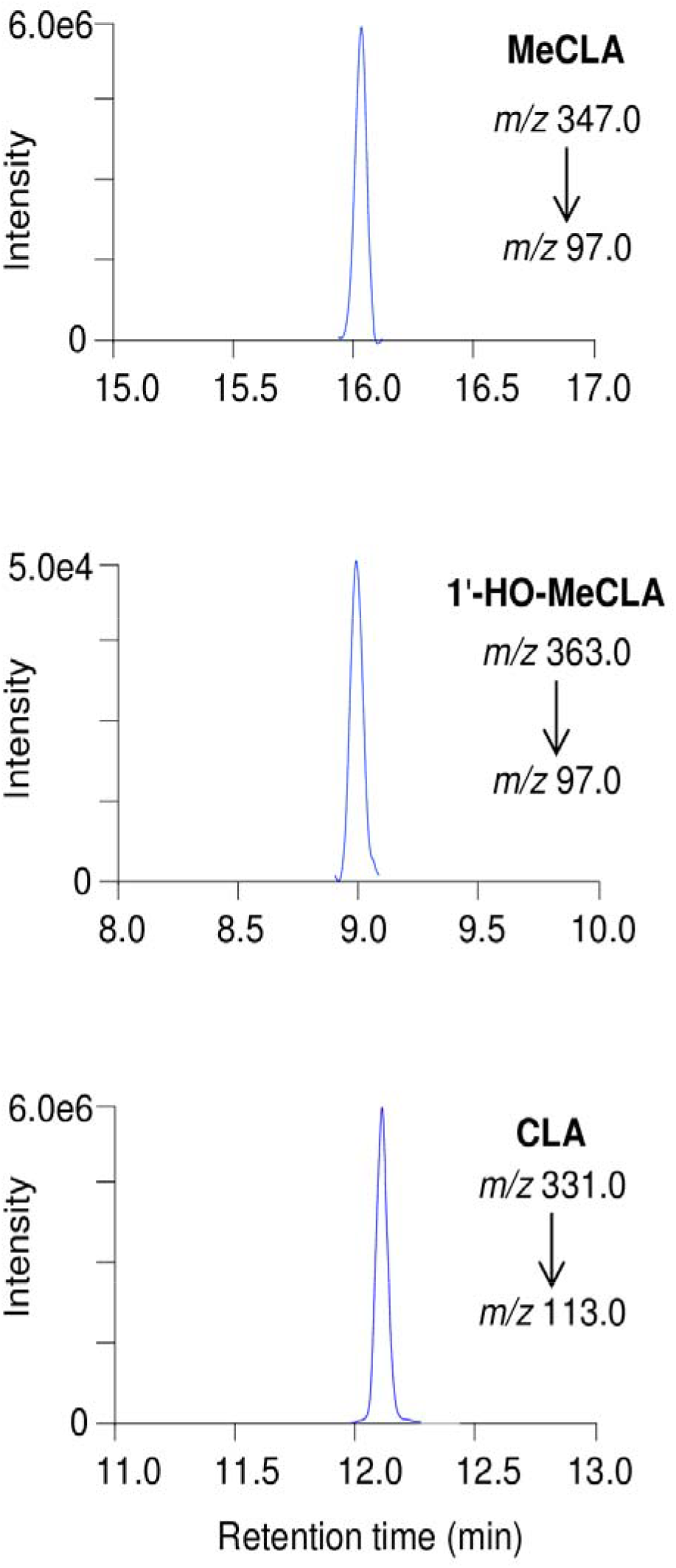
Most MeCLAs were converted to CLA. MeCLA was incubated with recombinant LBO proteins for 15 min. The extracts were analyzed by LC-MS/MS (MRM).

### 1’-HO-MeCLA is present in *atd14* mutants but not in *lbo* mutants

It is important to clarify if 1’-HO-MeCLA is an endogenous compound in plant tissues because there is a possibility that 1’-HO-MeCLA would only be produced in the heterologous expression system. Identification of 1’-HO-MeCLA was conducted using *atd14* mutant plants, because they lack a functional SL receptor and accumulate SLs due to negative feedback on the biosynthesis pathway. As a negative control, *lbo* mutant plants were also used. 1’-HO-MeCLA was detected from the basal part of shoots, and also root tissues of *atd14* mutants (Fig. 4). By contrast, CL and MeCLA, but not 1’-HO-MeCLA, were detected from both tissues of *lbo* mutants (Fig. 4, Brewer et al. 2016). These results clearly indicate that LBO may act to convert MeCLA into 1’-HO-MeCLA in plants.

**Figure 4.**
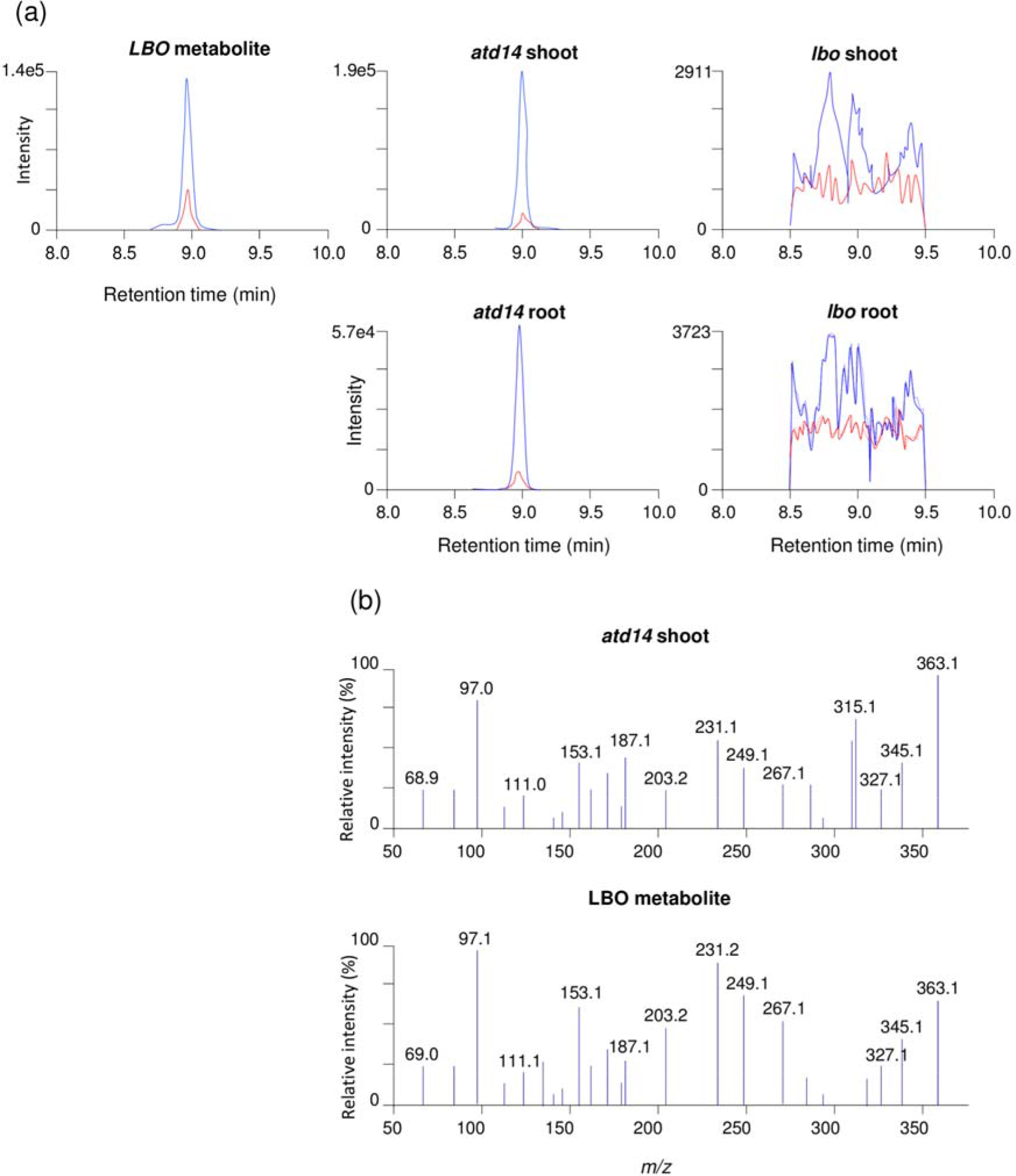
1’-HO-MeCLA was found from *atd14* shoot. Identification of endogenous 1’-HO-MeCLA in basal parts of shoot and root tissues was conducted. (a) MRM of chromatograms (363.0/97.0; *m/z* in positive mode) of *atd14* mutants (*Middle*) and *lbo* mutants (*Light*). (b) Product ion spectra derived from endogenous 1’-HO-MeCLA in basal parts of shoot of *atd14* mutants.

### Production of 1’-HO-MeCLA correlates with shoot branching

We previously described additional alleles of mutation in the *LBO* gene (Brewer et al. 2016). *lbo-2* plants have a point mutation in the predicted catalytic domain and display significant extra branching. *lbo-3* plants have a point mutation elsewhere in the gene and have a branching phenotype that is much weaker than *lbo-2* (Brewer et al., 2016). LBO-2 and LBO-3 proteins were produced in *E. coli* heterologous expression system and enzymatic activities to produce 1’-HO-MeCLA were examined. The very low conversion of MeCLA to 1’-HO-MeCLA by LBO-2 enzyme activity (Fig. 5) relates well to the mutant shoot branching phenotype. However, LBO-3 appears to have normal function in our assay (Fig. 5).

**Figure 5.**
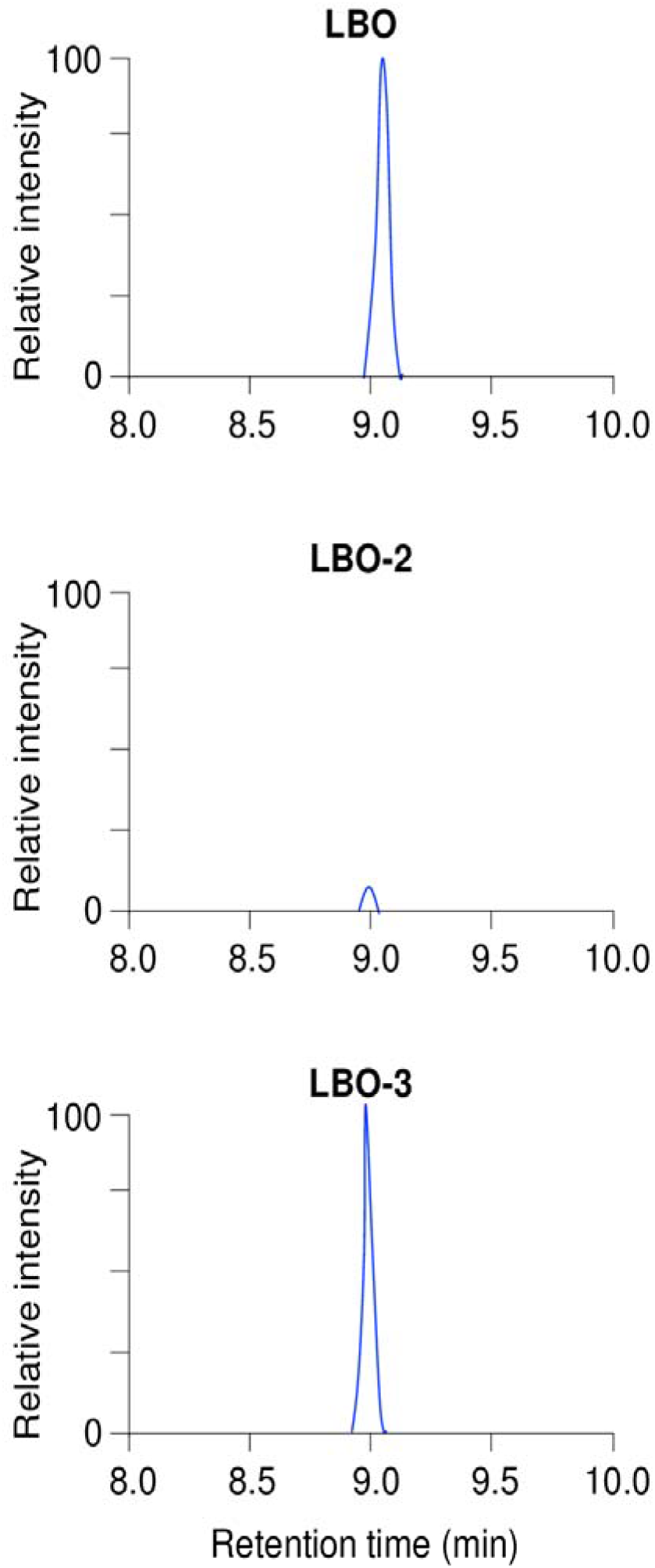
Production of 1’-HO-MeCLA is very low in LBO-2. MeCLA was incubated with each recombinant protein for 15 min and extracts were analyzed by LC-MS/MS. MRM chromatograms of 1’-HO-MeCLA (363.0/97.0; *m/z* in positive mode) are shown.

### Conversion of MeCLA into 1’-HO-MeCLA is conserved among different plant species

Tomato, maize, and sorghum have one LBO homolog each and their recombinant LBO proteins were expressed in *E. coli*. Not only *Arabidopsis* LBO but also the other LBO proteins examined converted MeCLA into 1’-HO-MeCLA (Fig. 6), where the major reaction product was CLA (Supplemental Fig. S1).

**Figure 6.**
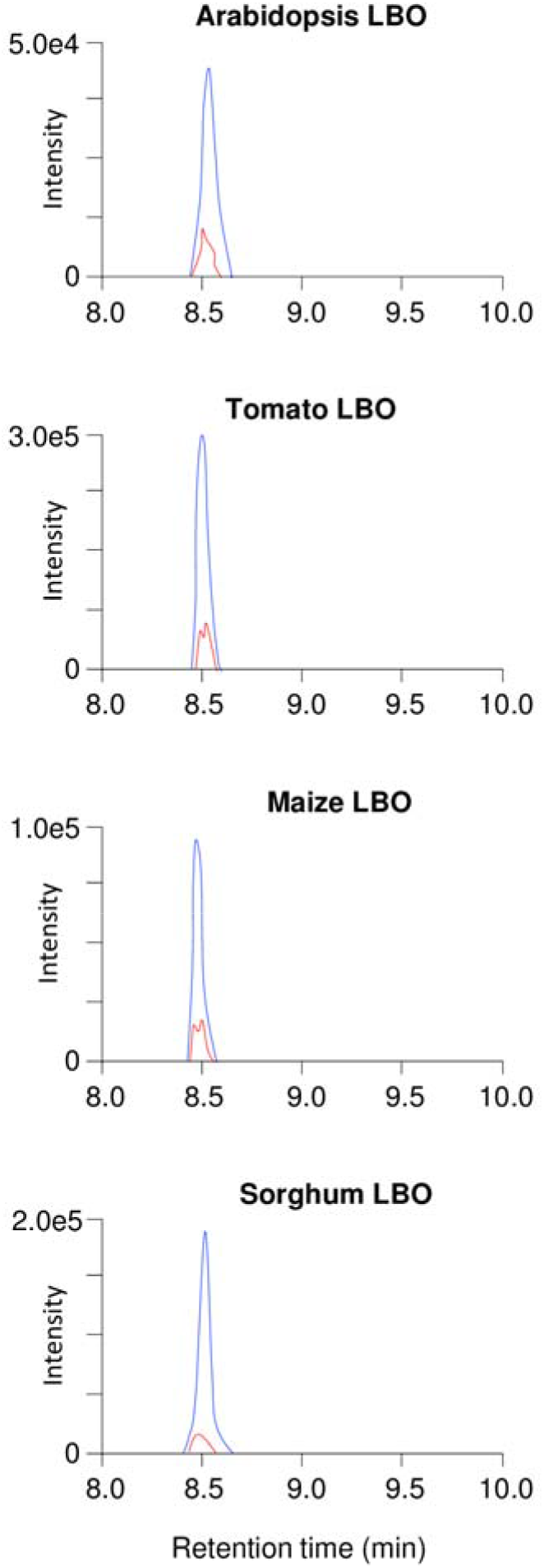
Conversion of MeCLA into 1’-HO-MeCLA is conserved among different plant species. MeCLA was incubated with each recombinant protein for 15 min and extracts were analyzed by LC-MS/MS. MRM chromatograms of 1’-HO-MeCLA (363.0/97.0; *m/z* in positive mode) are shown.

It is intriguing to test if LBO has an ability to produce canonical SLs or not. Tomato plants produce canonical SLs such as solanacol and orobanchol. Tomato MAX1 expressed in yeast cannot produce canonical SLs from CL (Yoneyama et al., 2018). Accordingly, there is a possibility that tomato LBO produces canonical SLs including solanacol and orobanchol. However, tomato LBO produced neither solanacol nor orobanchol from MeCLA (Data not shown). In addition, tomato LBO did not convert 4DO into solanacol or orobanchol, either (data not shown). Similar results were obtained with sorghum or maize LBOs. Sorghum LBO produced neither 5-deoxystrigol (5DS) nor sorgomol, two major canonical SLs of sorghum (cv Hybrid), from MeCLA. Although it was proposed that sorgomol is produced from 5DS (Motonami et al., 2013), LBO did not produce sorgomol from 5DS (data not shown). Maize plants produce zealactone (Charnikhova et al., 2017; Xie et al., 2017) and zeapyranolactone (Charnikhova et al., 2018), non-canonical SLs with unique structures. Maize LBO did not produce these SLs from MeCLA (Data not shown).

### Endogenous non-canonical SLs in *Arabidopsis*

CYP711A2, one of rice MAX1 homologs, produces 4-deoxyorobanchol (4DO) via 18-HO-CLA from CL (Yoneyama et al., 2018). This suggests that not only 1’-HO-MeCLA but also other HO-CL derivatives including HO-CLs, HO-CLAs, and HO-MeCLAs are endogenous compounds in *Arabidopsis,* and some of them may be substrates for MAX1 and LBO. Therefore, endogenous SLs in *atd14*, *max1* and *lbo* mutants were investigated in detail. Synthetic standards of 2-, 3-, 4-, 16- and 18-HO-CL (Fig. 7) were prepared and used for LC-MS/MS analyses. HO-CLAs (Fig. 7) were obtained by conversion of the corresponding HO-CLs by MAX1 expressed in yeast. HO-MeCLAs (Fig. 7) were obtained by methylation of the corresponding HO-CLAs with diazomethane.

**Figure 7.**
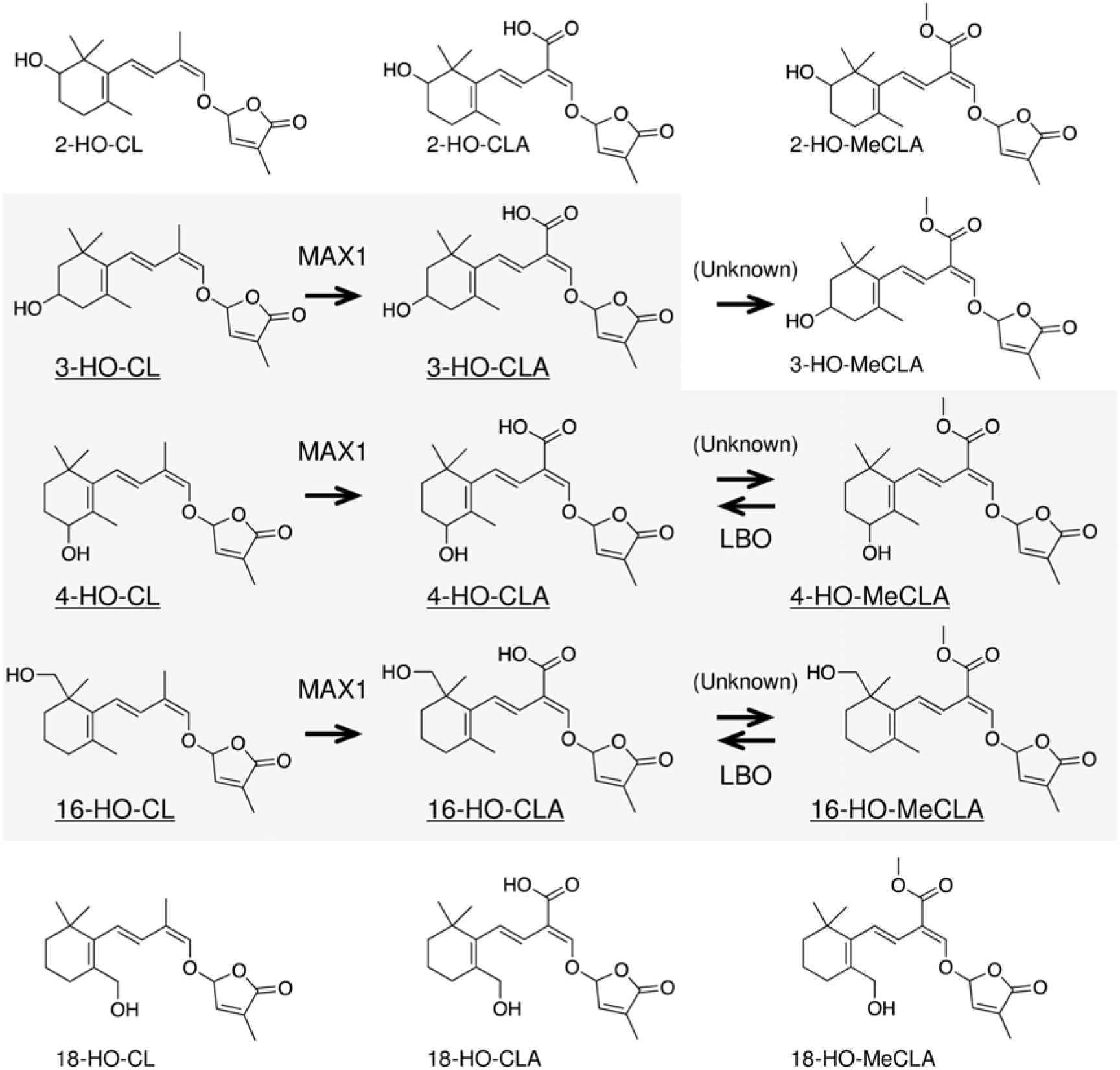
Structures of HO-CLs, HO-CLAs and HO-MeCLAs and a proposed strigolactone biosynthesis pathway in *Arabidopsis*. The present study shows that 3-, 4-, and 16-HO-CL derivatives are predominant and produced through MAX1 and LBO in *Arabidopsis.*

Basal parts of *Arabidopsis* shoot were harvested when the shoot branching phenotype was clearly observed (Supplemental Fig. S2). From *atd14* mutants, 3-, 4-, and 16-HO-CLs, 3-, 4-, and 16-HO-CLAs, and 4- and 16-HO-MeCLAs, in addition to CL, CLA, and MeCLA, were detected (Fig. 8).

3-, 4-, and 16-HO-CLs and CL were detected from basal parts of *max1* mutants (Fig. 8). Although 3-, 4-, and 16-HO-CLAs were detected, even from Col-0 plants (Supplemental Fig. S3), these HO-CLAs were not detected in *max1* mutants (Fig. 8). By comparing peak areas of MRM chromatograms between *atd14* and *max1* mutants (Fig. 8), 3-, 4-, and 16-HO-CLs appeared to accumulate in *max1* mutants.

**Figure 8.**
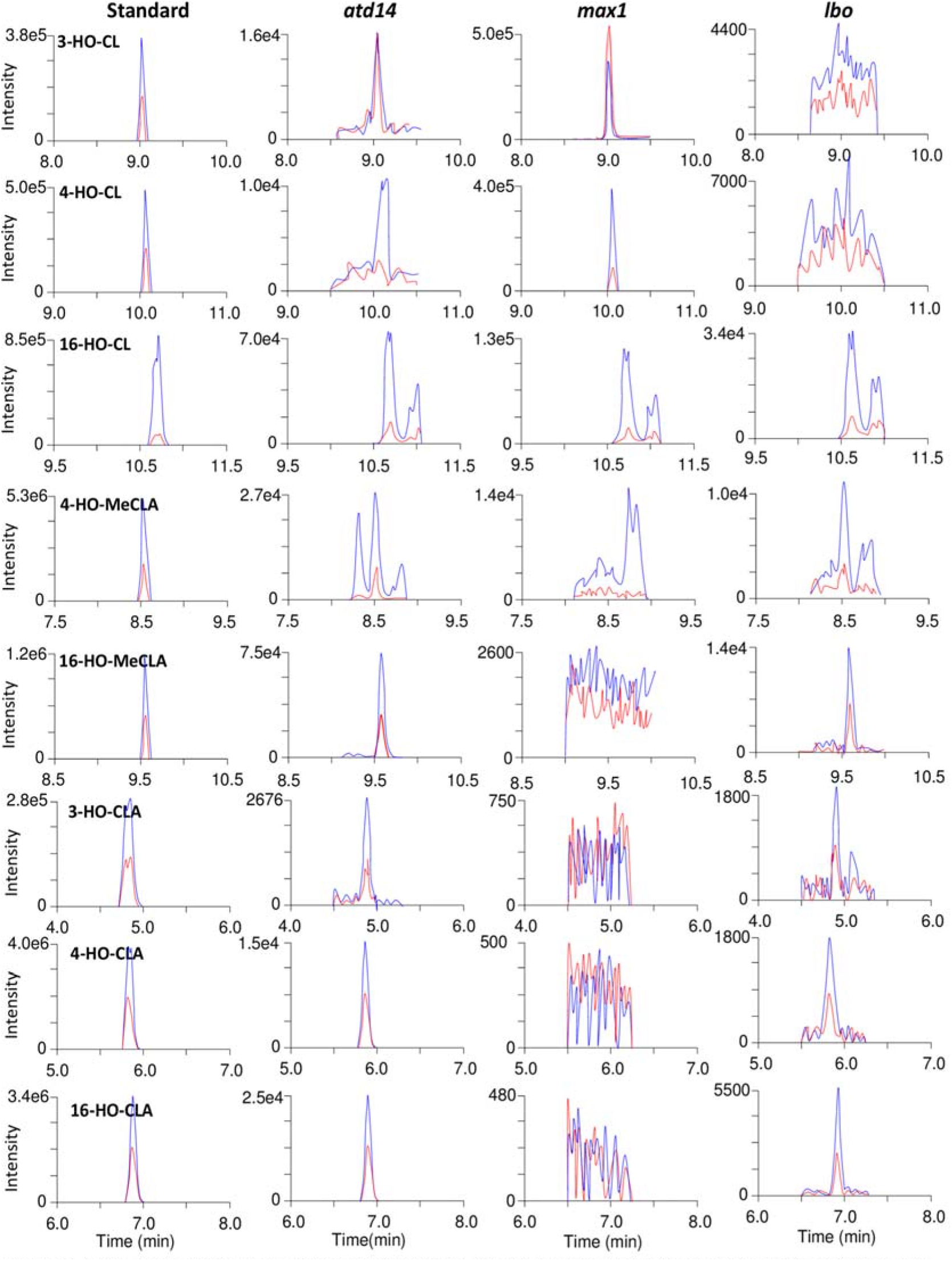
(a) HO-CLs and HO-MeCLAs are detected from ethyl acetate soluble fractions, and (b) HO-CLAs were from acidic fractions in basal parts of shoot tissues of *atd14* mutants, *max1* mutants, and *lbo* mutants. 3-, 4-, and 16-HO-CLs appeared to accumulate in *max1* mutants and 4-, and 16-HO-MeCLA in *lbo* mutants. MRM chromatograms of 3-HO-CL (blue, 301.0/97.0; red, 319.0/205.0; *m/z* in positive mode), 4-HO-CL (blue, 301.0/97.0; red, 301.0/148.0; *m/z* in positive mode), 16-HO-CL (blue, 301.0/97.0; red, 301.0/189.0; *m/z* in positive mode), 4-HO-MeCLA (blue, 345.0/97.0; red, 345.0/216.0; *m/z* in positive mode), 16-HO-MeCLA (blue, 345.0/97.0; red, 363.0/97.0; *m/z* in positive mode), 3-, 4-, and 16-HO-CLA (blue, 347.0/113.0; red, 347.0/69.0; *m/z* in negative mode) are shown.

### 4- and 16-HO-MeCLAs are potential substrates for LBO

In addition to CL, CLA, and MeCLA, 16-HO-CL, 3-, 4-, 16-HO-CLAs, 4- and 16-HO-MeCLAs were found in *lbo* mutants (Fig. 8). MeCLA was found to be a substrate for LBO (Brewer et al., 2016) and therefore these HO-MeCLAs also can be potential substrates for LBO.

Then, these HO-CL derivatives were incubated with recombinant LBO proteins as potential substrates. 4- and 16-HO-MeCLAs, but not other HO-CL derivatives were consumed by LBO. Although we searched for LBO products of 4- and 16-HO-MeCLAs with the D-ring fragment (*m/z* 97) as an indicator by LC-MS/MS, we could not find any candidates for LBO products (Fig. 9). As in the case of MeCLA, the corresponding HO-CLA was detected as a major reaction product.

**Figure 9.**
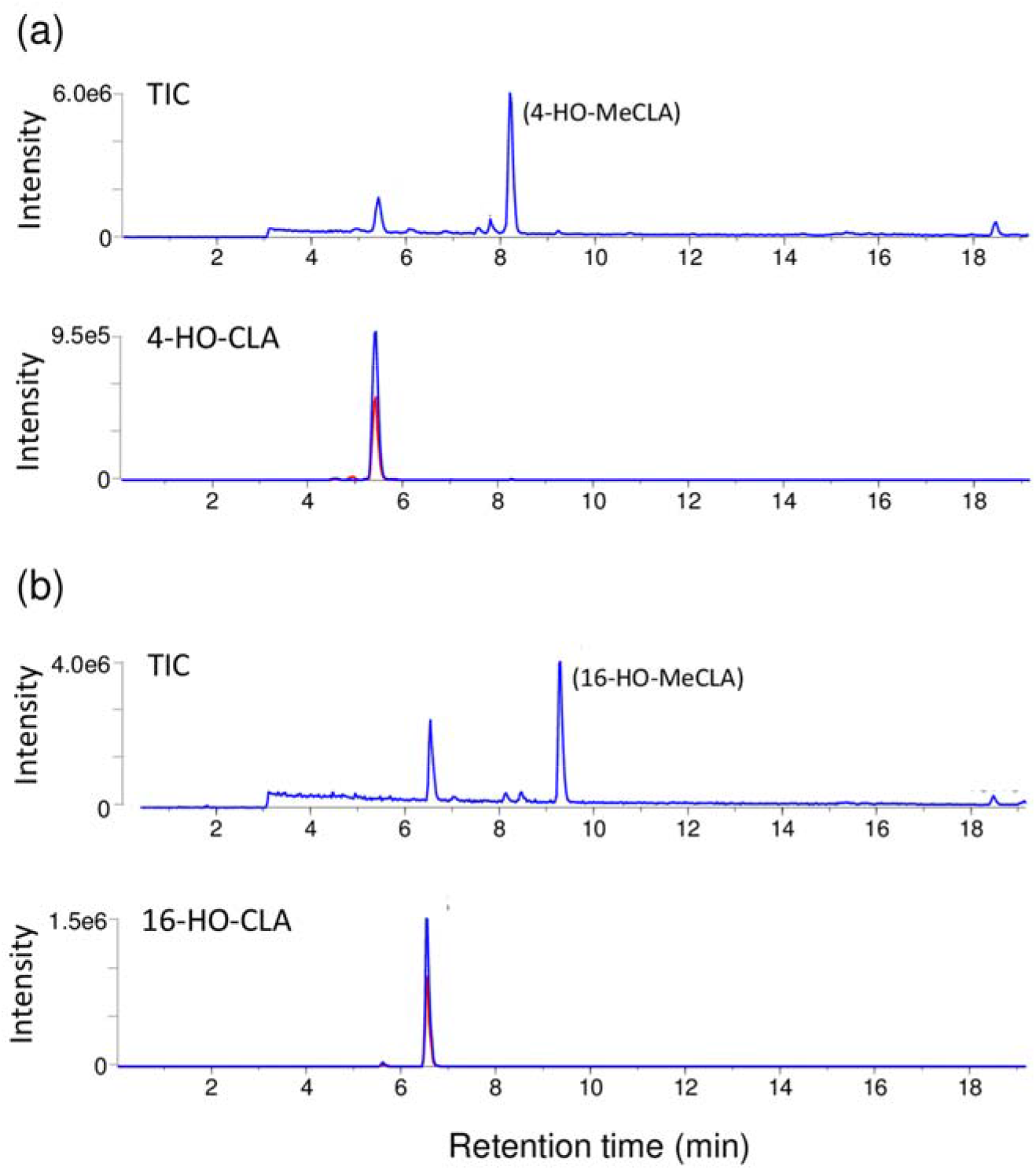
Recombinant LBO proteins convert 4-, and 16-HO-MeCLA mainly into the corresponding HO-CLAs. Each substrate was incubated for 15 min. The extracts were analyzed by LC-MS/MS with the D-ring fragment (*m/z* 97) as an indicator to identify the metabolites from each HO-CLA-fed LBO. Total Ion and MRM chromatograms of HO-CLAs (blue, 347.0/113.0; red, 347.0/69.0; *m/z* in negative mode) are shown.

## DISCUSSION

The present study demonstrated that the structure of [MeCLA+16] Da is 1’-HO-MeCLA and this LBO metabolite is endogenous in *Arabidopsis* tissues. 1’-HO-MeCLA was also produced by MeCLA-fed maize, tomato and sorghum LBO proteins, suggesting that this conversion of MeCLA into 1’-HO-MeCLA is highly conserved among different seed plant species.

Then the question arises whether 1’-HO-MeCLA is a strong shoot branching inhibitor or not. So far, 1’-HO-MeCLA has not been examined for its effect on shoot branching. Unfortunately, at this present time, the synthetic standard for 1’-HO-MeCLA is not available. As mentioned, the yield of 1’-HO-MeCLA by LBO protein reaction is too low to obtain enough for shoot branching assays. Since the substitution of 1’-HO-MeCLA is very unstable and could be readily converted, it is possible that 1’-HO-MeCLA is a precursor for an unknown, downstream shoot branching inhibitor(s) and a subsequent unknown enzyme(s) converts 1’-HO-MeCLA into the true shoot branching inhibitor(s). However, we cannot yet find any candidate compounds that are likely to be derived from 1’-HO-MeCLA from *atd14* mutants (data not shown). *LBO* was uncovered from transcriptomics (Brewer et al. 2016) and similar methods recently led to the discovery that CYP722C from cowpea and tomato converts CLA directly to orobanchol (Wakabayashi et al. 2019), and that a 2-oxoglutarate dependent dioxygenase (2-OGD) from a nearby clade to *LBO* is involved in SL biosynthesis in *Lotus japonicus* (Mori et al. 2020). We will continue reverse genetic and mass spectrometric approaches to find related SL biosynthetic genes and shoot branching inhibitors. CLA seems to be a key precursor for canonical SLs. We will test how LBO relates to CLA and canonical SLs by identifying *lbo* mutants from plants that produce canonical SLs.

Worthy of attention here is that the LBO protein assay produces much more CLA from MeCLA than 1’-HO-MeCLA. Such *O*-demethylations have been reported for 2-OGDs, thebaine 6-*O*-demethylase and codeine *O*-demethylase, catalyzing *O*-demethylation in the final steps of morphine biosynthesis (Hagel and Facchini, 2010). Therefore, we cannot exclude the possibility that the main function of LBO is demethylation of MeCLA and that 1’-HO-MeCLA is just an intermediate for demethylation. This is somewhat difficult to reconcile with our previous result that showed that CLA was detected from *lbo* mutants, but not from its wild type (Brewer et al., 2016), indicating that CLA accumulates in *lbo* mutants. As methyltransferase is proposed to be involved in conversion from CLA into MeCLA and should be functional in *lbo* mutants. Thus, CLA would not be expected to accumulate in *lbo* mutants unless the methyltransferase was somehow downregulated. Identification of the methyltransferase will clarify the meaning of methylation and demethylation in the production of shoot branching inhibitors.

*lbo-2* mutants with a point mutation in the predicted catalytic domain display extra shoot branching. The present study demonstrated that recombinant LBO-2 protein is very weak at converting MeCLA into 1’-HO-MeCLA (Fig. 5). In contrast, LBO-3 appears to have normal function in our assay (Fig. 5). The shoot branching phenotype of *lbo-3* mutants with a point mutation elsewhere in the gene is much weaker than *lbo-2,* and only just significantly more than wild type (Brewer et al., 2016). It is possible that the protocol was not sensitive enough to observe subtle defects in reaction efficiency. Alternatively, the LBO-3 mutation may reveal an unknown protein functional or interaction domain at the mutation site, which only affects its bioactivity *in planta*. It may be useful to test LBO and LBO-3 in combination with other SL biosynthesis enzymes as they become discovered.

In addition to 1’-HO-MeCLA, other unstable non-canonical SLs were found in the basal parts of shoot tissues (Fig. 8). So far, identification of SLs has been mainly conducted from root tissues and this is the first report to show that SLs exist in basal parts of shoots of *Arabidopsis*. There were no apparent differences in SL levels between the two tissues when peak areas were compared (Fig. 4). In contrast, levels of SLs are very low or undetectable from shoot of sorghum (Yoneyama et al., 2007) and rice plants (Umehara et al., 2010). *Arabidopsis* could be quite particular in containing the same levels of SLs in the basal part of shoot and root tissues. Perhaps this is because *Arabidopsis* is a non-host of AM fungi. Even though *Arabidopsis* was reported to produce orobanchol (Goldwasser et al., 2008; Kohen et al., 2011), a canonical SL that is widely distributed in plant kingdom (Yoneyama et al., 2008; Yoneyama et al., 2011), we could detect neither orobanchol nor any other known canonical SLs from *Arabidopsis* tissues (data not shown). Non-canonical SLs seem to be predominant in *Arabidopsis* and may not be released into the soil because *Arabidopsis* does not need to attract AM fungi to form a relationship with them. However, there are hints that SLs in *Arabidopsis* may promote interaction with other beneficial soil fungi (Carvalhais et al. 2019). So, there is likely much more to learn on that topic.

As summarized in Fig 7, MAX1 oxidizes C-19 methyl group to carboxylic acid not only in CL, but also in HO-CLs in *Arabidopsis* plants. Baz et al. (2018) also detected 3-HO-CL from rice *d14* mutant roots and demonstrated that 9-*cis*-3-HO-β-apo-10’-carotenal-fed to OsCCD8 is converted into 3-HO-CL. These results suggest that HO-CLs are also converted by MAX3 from HO-carotenal. The *Arabidopsis* MAX1 enzyme has the ability to convert 2-HO-CL and 18-HO-CL into respective HO-CLAs. However, these HO-CL derivatives could not be found from *Arabidopsis* plants. It is intriguing why *Arabidopsis* produces such various and particular HO-CL derivatives.

## CONCLUSION

Deciphering the whole SL biosynthetic pathway and characterization of yet unidentified biosynthetic intermediates is essential for devising new strategies to regulate the multiple functions of SLs through manipulation of SL production and exudation, both quantitatively and qualitatively. It should be noted that SL production and exudation vary with plant species (even between cultivars or genotypes of the same plant species), growth conditions, and growth stages. In the present study, we have unveiled the enzymatic functions of LBO and MAX1 and their substrates and products downstream of CL in the SL biosynthetic pathway in *Arabidopsis*. As most seed plant species sequenced so far contain a single *LBO* gene, and the *LBO* gene lineage appears to have been derived deep in plant evolutionary history (Walker et al., 2019), the biological function of LBO is likely to be highly conserved in the plant kingdom.

## Materials and Methods

### Plant material

The *lbo-1* and *max1-4* were from our *Arabidopsis* laboratory stocks (Brewer et al., 2016) and the *atd14-2* mutant was obtained from a TILLING project in the Columbia-0 (Col-0) ecotype. To extract total RNAs, tomato (cv Ailisa Craig; Nomura et al., 2005), maize (cv B73; Yoneyama et al., 2018) and sorghum (cv Hybrid; Yoneyama et al., 2008) were used.

### Chemicals

3-, 4-, and 18-HO-CLs were synthesized as described previously (Mori et al. 2016; Baz et al. 2018). 2- and 16-HO-CLs were synthesized using the same strategy as the synthesis of 3-and 18-HO-CLs (Baz et al., 2018; Mori et al., 2016). The detailed synthesis will be published elsewhere. 2-, 3-, 4-, 16- and 18-HO-CLA were obtained by MAX1 microsome assay using the corresponding HO-CLs. For this, MAX1 expressed in yeast (*Saccharomyces cerevisiae*) was prepared as described previously (Abe et al., 2014, Yoneyama et al., 2018). 2-, 3-, 4-, 16- and 18-HO-MeCLA were prepared by methylation of the corresponding HO-CLAs with diazomethane.

### Synthesis of methyl-*d*_3_ carlactonoate (1’-*d*_3_-MeCLA) (Scheme S1)

(*E*)-4-(2,6,6-Trimethylcyclohex-1-en-1-yl) but-3-enoic acid was synthesized as reported (Abe et al., 2014). To a solution of the C_13_-carboxylic acid (88.1 mg, 0.42 mmol) in acetone (2 mL), K_2_CO_3_ (174 mg, 1.26 mmol) and methyl-*d_3_* iodide (305 mg, 131 μL, 2.1 mmol) were added. The mixture was stirred at room temperature for 21 h under argon. After being concentrated under nitrogen gas flow, the residue was dissolved with ether and water. The organic phase was washed with water and dried over MgSO_4_. Filtration and evaporation of the solvent afforded C_13_-carboxylic acid methyl-*d_3_* ester (82.3 mg, 0.37 mmol, 87%), which was pure enough for the next reaction. Ester condensation of the methyl-*d_3_* ester (82.3 mg, 0.37 mmol) with ethyl formate (98 mg, 106 μL, 1.32 mmol) by the use of sodium hydride (13.3 mg, 0.56 mmol) in *N,N*-dimethylformamide (1 mL) followed by alkylation with racemic 4-bromo-2-methyl-2-buten-4-olide (99 mg, 55 μL, 0.56 mmol) (Abe et al., 2014) provided 1’-*d*_3_-MeCLA and ethyl carlactonoate (EtCLA, a transesterification product). Purification by silica gel column chromatography (Kieselgel 60, Merck, *n*-hexane-ethyl acetate stepwise) and semi-preparative HPLC (Inertsil SIL-100A, GL Sciences, 5% ethanol in *n*-hexane) gave 1’-*d*_3_-MeCLA (2.5 mg, 0.0072mmol, 1.9%). **1’-*d*_3_-MeCLA**: HR-ESI-TOF-MS *m/z*: 372.1855 [M+Na]^+^ (calcd. for C_20_H_23_D_3_NaO_5_^+^, *m/z*: 372.1861).

### Synthesis of methyl 18-*d*_3_-carlactonoate (18-*d*_3_-MeCLA) (Scheme S2)

6,6-Dimethyl-2-(methyl-*d_3_*)cyclohex-1-en-1-yl trifluoromethanesulfonate was synthesized as reported (Tanaka et al., 2007). A mixture of the triflate (5.30 g, 19.3 mmol), triethylamine (7.80 g, 10.7 mL, 77.2 mmol), methyl 3-butenoate (3.86 g, 4.11 mL, 38.6 mmol), and bis(triphenylphosphine)palladium(II) dichloride (1.35 g, 1.92 mmol) in *N,N*-dimethylformamide (50 mL) was stirred at 100°C for 17 h under argon. The reaction mixture was cooled, quenched by pouring into 1 N HCl, and extracted with ether. The organic phase was washed with brine and water, dried over MgSO_4_, and concentrated in vacuo. Purification by silica gel column chromatography (Kieselgel 60, Merck, *n*-hexane-ether stepwise) gave crude methyl (*E*)-4-(6,6-dimethyl-2-(methyl-*d_3_*)cyclohex-1-en-1-yl)but-3-enoate (1.31 g, 5.8 mmol, 30%), which was used for the next reaction without further purification. Ester condensation of the deuterium-labeled ester (108 mg, 0.48 mmol) with methyl formate (86.4 mg, 89 μL, 1.44 mmol) by the use of sodium hydride (11.5 mg, 0.48 mmol) in *N,N*-dimethylformamide (1 mL) followed by alkylation with racemic 4-bromo-2-methyl-2-buten-4-olide (85 mg, 47 μL, 0.48 mmol) (Abe et al., 2014) provided 18-*d*_3_-MeCLA. Purification by silica gel column chromatography (Kieselgel 60, Merck, *n*-hexane-ethyl acetate stepwise), semi-preparative normal-phase HPLC (Inertsil SIL-100A, GL Sciences, 5% ethanol in *n*-hexane) and semi-preparative reversed-phase HPLC (InertSustain C18, GL Sciences, 85% acetonitrile in water) gave 18-*d*_3_-MeCLA (1.7 mg, 0.0049 mmol, 1.0%). **18-*d*_3_-MeCLA**: HR-ESI-TOF-MS *m/z*: 350.2058 [M + H]^+^ (calcd. for C_20_H_24_D_3_O_5_^+^, *m/z*: 350.2041).

### Cloning

The primer sequences used are listed in Supporting Information TableS1. Total RNAs were extracted from the shoots and roots of plant materials using an RNeasy Plant Mini Kit (Qiagen, Hilden, Germany) and employed to synthesize single-strand cDNAs by a SuperScript III First-Strand Synthesis System (Invitrogen, Waltham, MA, USA). PCR amplification was performed using PrimeSTAR HS DNA polymerase (TAKARA Bio Inc., Kusatsu, Japan) with/without GC buffer for accurate amplification of GC rich targets. The full-length cDNAs were cloned into the pENTR vector and then transferred to pET300 vector by the Gateway system (Invitrogen). Recombinant plasmid DNA was transferred to *Escherichia coli* strain Rosetta 2(DE3)pLysS (Novagen). At least four colonies for each experiment were sequenced to check for errors in the PCR. Sequence alignment was performed using MAC VECTOR software (Mac Vector Inc., Apex, NC, USA).

### Heterologous expression in *E. coli*

Heterologous expression of LBO in *E. coli* was carried out as described previously (Brewer et al., 2016). Briefly, transformed colonies were grown in LB media (0.5% yeast extract, 1% Bacto Tryptone, 1% NaCl) with carbenicilin (100 μg/mL) at 37 °C in a shaking incubator (180 rpm) until the cell density reached an OD_600_ of 0.5-0.8. After isopropyl-D-1-thiogalactopyranoside (1 mM) was added, transformed *E. coli* were incubated at 20 °C for 14-16 h. To prepare enzyme fractions, *E. coli* cells were collected by centrifugation of 10,000 x *g* for 1 min and suspended in 20 mM phosphate buffer (pH 7.4). The suspend cells were mechanically lysed by using a high-pressure homogenizer (Emulsi Flex B15; AVESTIN) and then, centrifuged at 15,000 x *g* for 5 min at 4°C.

### LBO enzyme assays and metabolite extraction

Crude protein fraction (5 mL) was incubated with 4 mM 2-oxoglutarate, 0.5 mM iron ascorbate, 5 mM ascorbic acid, and 12.5 μg of test substrates at 27°C for 20 min, similar to the previous report (Brewer et al., 2016). The reaction mixture was extracted with 5 mL ethyl acetate twice. The ethyl acetate soluble fraction was dried with sodium sulfate and evaporated under nitrogen gas flow at 40°C with care not to completely dry. Crude extract samples were kept at −20°C until LC-MS analysis.

### SL identification in *A. thaliana*

*Arabidopsis* seeds were sterilized in 1% sodium hypochlorite solution for 10 min and rinsed with sterile water. Seeds were sown on agar (0.5% gellangum with 1/2 Murashige and Skoog medium and 1% sucrose), stratified at 4°C for 2 days, and grown for 10 days under a photoperiod, 14 h: 10 h, light (150 mol m^−2^ s^−1^): dark, at room temperature. Then, healthy and uniform seedlings were transplanted on soils [horticultural soil: vermiculite = 1: 2 (v/v)] and further grown until branching phenotype became clear. Basal parts of shoot tissues were harvested, extracted with ethyl acetate for at least 2 days and crude extracts were purified by DEA and silica Sep-pack cartridge as reported previously (Brewer et al., 2016).

### LC-MS/MS analysis

SLs were analyzed by LC-MS/MS as reported previously (Abe et al., 2014). Briefly, LC-MS/MS analysis (MRM, multiple reaction monitoring and PIS, product ion scan) of proton adduct ions was performed with a triple quadruple/linear ion trap instrument (QTRAP5500; AB Sciex, Old Connecticut Path Framingham, MA, USA) with an electrospray source. HPLC separation was performed on a UHPLC (Nexera X2; Shimadzu) equipped with an ODS column (Kinetex C18, 2.1 × 150 mm, 1.7 m; Phenomenex) with a linear gradient of 35% acetonitrile (0 min) to 95% acetonitrile (20 min). The column oven temperature was maintained at 30°C.

## Supplemental Data

**Supplemental Figure S1**. Detection of CLA from recombinant LBO proteins of various plants.

**Supplemental Figure S2**. The number of shoot branching in *max1*, *lbo* and *atd14* mutants.

**Supplemental Figure S3**. Identification of HO-CLAs from Col-0.

## ACKNOWLEDGEMENTS

We would like to thank Nozomi Nanai for technical assistance. This study was supported by the Japan Science and Technology Research Promotion Program for Agriculture, Forestry, Fisheries and Food Industry, the Japan Society for the Promotion of Sciences (KAKENHI 15K07093, 16K07618, 16K18560) and the Japan Science and Technology Agency PRESTO (JPMJPR17QA).

